# Neural source dynamics of brain responses to continuous stimuli: speech processing from acoustics to comprehension

**DOI:** 10.1101/182881

**Authors:** Christian Brodbeck, Alessandro Presacco, Jonathan Z. Simon

## Abstract

Human experience often involves continuous sensory information that unfolds over time. This is true in particular for speech comprehension, where continuous acoustic signals are processed over seconds or even minutes. We show that brain responses to such continuous stimuli can be investigated in detail, for magnetoencephalography (MEG) data by combining linear kernel estimation with minimum norm source localization. Previous research has shown that the requirement to average data over many trials can be overcome by modeling the brain response as a linear convolution of the stimulus and a kernel, or response function, and estimating a kernel that predicts the response from the stimulus. However, such analysis has been typically restricted to sensor space. Here we demonstrate that this analysis can also be performed in neural source space. We first computed distributed minimum norm current source estimates for continuous MEG recordings, and then computed response functions for the current estimate at each source element, using the boosting algorithm with cross-validation. Permutation tests can then assess the significance of individual predictor variables as well as features of the corresponding spatio-temporal response functions. We demonstrate the viability of this technique by computing spatio-temporal response functions for speech stimuli, using predictor variables reflecting acoustic, lexical and semantic processing. Results indicate that processes related to comprehension of continuous speech can be differentiated anatomically as well as temporally: acoustic information engaged auditory cortex at short latencies, followed by responses over the central sulcus and inferior frontal gyrus, possibly related to somatosensory/motor cortex involvement in speech perception; lexical frequency was associated with a left-lateralized response in auditory cortex and subsequent bilateral frontal activity; and semantic composition was associated with bilateral temporal and frontal brain activity. We conclude that this technique can be used to study the neural processing of continuous stimuli in time and anatomical space with the millisecond temporal resolution of MEG. This suggests new avenues for analyzing neural processing of naturalistic stimuli, without the necessity of averaging over artificially short or truncated stimuli.

## 1 Introduction

In a natural environment, the brain frequently processes information in a continuous fashion. For example, when listening to continuous speech, information is extracted incrementally from an uninterrupted acoustic signal at multiple levels: phonetically relevant sound patterns are recognized and grouped into words, which in turn are integrated into phrases which are meaningful in the context of a larger discourse (e.g. Gaskell & Mirkovic, 2016). Contrary to this continuous mode of functioning, neuroimaging experiments typically isolate phenomena of interest with short, repetitive trials (for many examples, see e.g. Gazzaniga, Ivry, & Mangun, 2009). While such research unquestionably leads to valuable results, the lack of naturalness of the stimuli is associated with uncertainty of how generalizable such results are to real world settings (see e.g. Brennan, 2016). Consequently, there is a need for complementary research with more naturalistic stimuli.

Brain responses to continuous speech have been studied with functional magnetic resonance imaging (fMRI) (Brennan et al., 2012; Brennan, Stabler, Van Wagenen, Luh, & Hale, 2016; Chow et al., 2014; Willems, Frank, Nijhof, Hagoort, & van den Bosch, 2016). Hemodynamic changes have been shown to track inherent properties of words, such as word frequency, as well as properties of words in context, such as contextual probability of encountering a given word. However, the low temporal resolution of fMRI, typically sampled at or below 1 Hz, imposes several limitations on the phenomena that can be modeled. While the studies cited above suggest that the resolution is adequate to model responses with a timescale of individual words, this is not the case for processes at faster timescales such as phonetic perception, where relevant events last only tens of milliseconds. In addition, fMRI responses can be modeled in terms of brain regions which are or are not sensitive to a given variable, but the relative and absolute timing of different components of the response remain obscure. Thus, even when word-based variables are analyzed, hemodynamic responses are modeled as instantaneous effects of the relevant variable, convolved with the hemodynamic response function, but without taking into account the temporal relationship between the stimulus and different components of the brain response (e.g. Brennan et al., 2016; Willems et al., 2016).

In contrast to fMRI, electroencephalography (EEG) and magnetoencephalography (MEG) have the temporal resolution to track continuous processing with millisecond accuracy. Previous research has established that the dependency of the MEG or EEG response on a continuous stimulus variable can be modeled as a linear time-invariant system (Lalor, Pearlmutter, Reilly, McDarby, & Foxe, 2006). This technique has been originally developed for relating neurons' spiking behavior to continuous sensory stimuli (see Ringach & Shapley, 2004), but can be extended to MEG/EEG signals by modeling the response as a linear convolution of a stimulus variable with an impulse response function (see Figure 1). Given a known stimulus and a measured response, one can then estimate the optimal response function to predict the measured response from the stimulus. This technique has been used to model EEG responses to continuously changing visual stimuli, by modeling continuous EEG signals as the convolution of moment-by-moment stimulus luminance with an appropriate response function (Lalor et al., 2006). An analogous procedure has been used to estimate responses to amplitude modulated tones and noise (Lalor, Power, Reilly, & Foxe, 2009). As an extension of this procedure, the response to continuous speech has been modeled as a response to the level of momentary acoustic power, the acoustic envelope (Lalor & Foxe, 2010).

**Figure 1:**
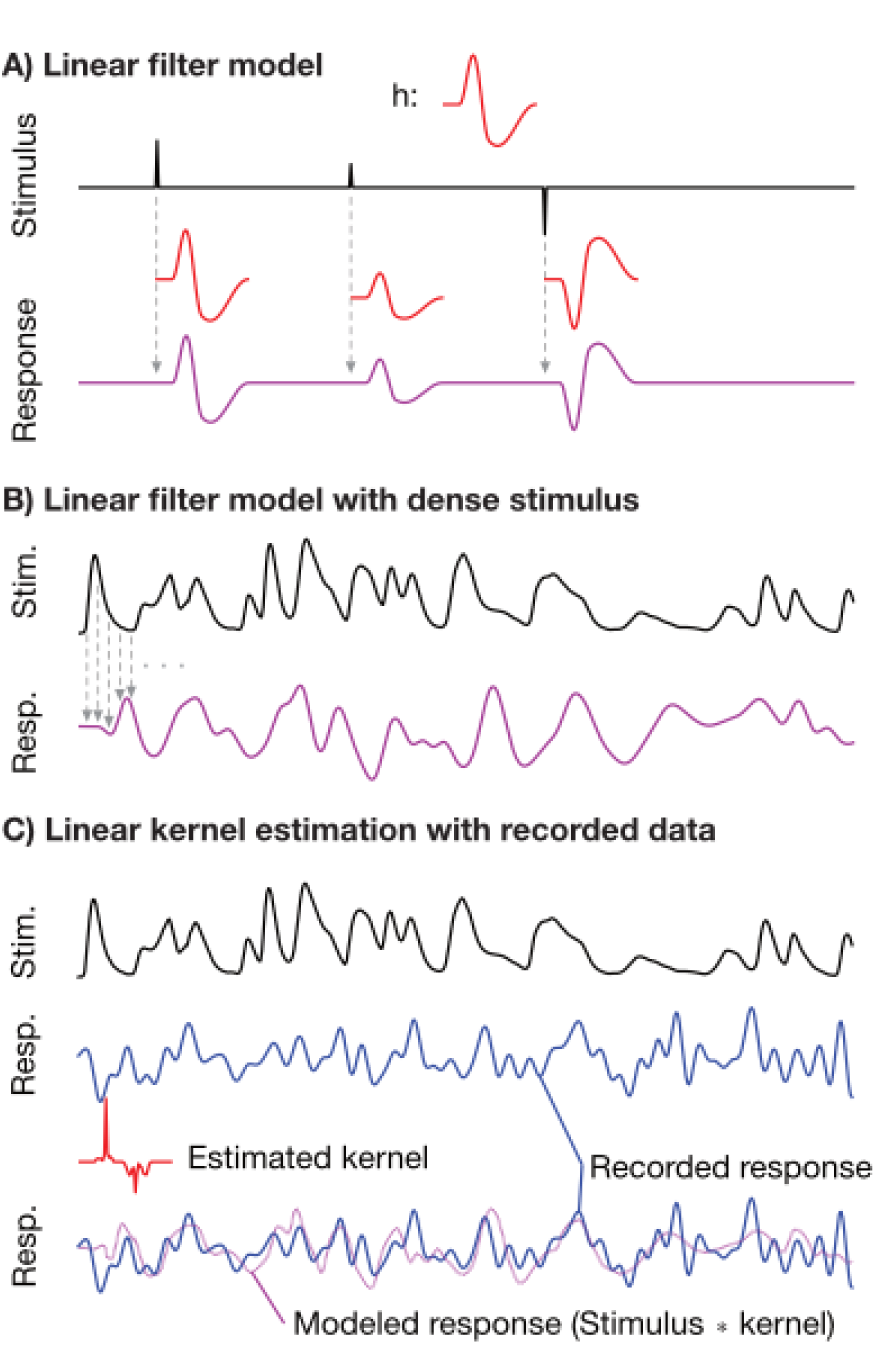
Linear filter model. The linear filter model *r* = *h* * *s* assumes that the response r is the convolution of the stimulus s with a response function, or kernel h. A) Illustrates the linear filter model for a simple stimulus with three discrete impulses. Since the impulses are spaced far apart relative to the size of the kernel, the shape of the kernel is apparent in the response. B) If the stimulus varies continuously, the convolution leads to a response in which the kernel is not discernible by eye. The response shown is obtained by convolving the stimulus with the same kernel as in A. C) If the stimulus as well as the response are known, different methods exist to estimate a kernel that optimally predicts the response given the stimulus. In the illustration, the simulated response is obtained by convolving the stimulus with the kernel shown under A and adding noise. The kernel is then estimated from the stimulus and the simulated response using boosting (see Methods). The modeled response, i.e. the stimulus convolved with the estimated kernel, can be compared to the actual response to determine the explanatory power of the model.

While the original formulation focused on purely sensory neurons, i.e. neurons whose response is a linear function of sensory input (Ringach & Shapley, 2004), the same method has also been applied successfully to determine cognitive influences on sensory processing. This can be achieved by modeling the signal as a response to a continuous predictor variable that represents a specific property of interest of the input stimulus. Thus, besides the acoustic envelope, the EEG response to continuous speech has been shown to reflect categorical representations of phonemes (Di Liberto, O’Sullivan, & Lalor, 2015). Furthermore, using stimuli in which speech from multiple talkers is mixed, it has been shown that the response function to the acoustic envelope can be divided into an earlier component around 50 ms that responds to the acoustic power in the overall stimulus, and a later component around 100 ms that responds to the acoustic envelope of the attended speech stream but not the unattended one (Ding & Simon, 2012b, 2012a).

While this research shows that response functions for continuous stimuli can be estimated, and that they can track not just sensory but also cognitive processes, all the above studies estimated response functions using only sensor space data. Topographic distributions of response functions have been assessed using equivalent current dipole localization (Lalor et al., 2009; Ding & Simon, 2012a) but this does not use the full localizing power of MEG. For investigating cognitive processing of sensory signals in particular, better source localization has the potential to separate response functions to different stimulus properties through anatomical separation of the brain response. In this paper, we propose to use distributed minimum norm source estimates to localize MEG data before estimating response functions. We developed a procedure in which source estimates are computed for continuous raw data, response functions are estimated independently at each virtual current dipole of the source model, and these individual response functions are then recombined to create a picture of the brain's responses to different functional aspects of the continuous stimulus, in both time and anatomical space. In other words, source localization is used to decompose the raw signal based on likely anatomical origin, and this decomposition is then used to estimate each potential source location's response to a particular stimulus variable.

To test and demonstrate this procedure, we analyzed data from participants listening to segments of a narrated story. We show that 6 minutes of data per participant is enough to estimate response functions that are reliable across subjects. In order to demonstrate the ability to localize responses in different brain regions, we focused on predictor variables with clearly different predictions for their anatomical localization and temporal response characteristics (see Figure 2): the response to the acoustic envelope of the speech signal should be associated with at least two strong components around 50 and 100 ms latency, in auditory cortex; previous studies suggest that the latter component is posterior to the former (Ding & Simon, 2012a). Responses associated with word recognition were assessed via lexical frequency, which is known to be one of the strongest predictors of lexical processing in general (see e.g. Baayen, Milin, & Ramscar, 2016). Higher frequency is associated with faster recognition of spoken words (e.g. Connine, Mullennix, Shernoff, & Yelen, 1990; Meunier & Segui, 1999; Dahan, Magnuson, & Tanenhaus, 2001) and is associated with lower amplitudes in event related potentials to single spoken words (Dufour, Brunellière, & Frauenfelder, 2013). FMRI investigations indicate a corresponding reduction in left-hemispheric temporal and frontal activity when processing more frequent compared to less frequent words in a narrated story (Brennan et al., 2016). Responses associated with higher levels of language processing beyond word recognition were assessed with an estimate of the amount of semantic combinatory processing over the course of the speech stimulus. This estimate was based on the presence of constructions associated with semantic composition operations, which previous MEG studies localized to the anterior temporal lobe (Bemis & Pylkkänen, 2011, 2012; Westerlund, Kästner, AI Kaabi, & Pylkkänen, 2015). This variable is relatively coarse and likely to be correlated with other variables reflecting structural integration, such as constituent size, associated with left temporal and inferior frontal activity (e.g., Pallier, Devauchelle, & Dehaene, 2011; Brennan et al., 2012). Consequently, this variable was treated as a rough estimate of multi-word integration processes during story comprehension, likely to be associated with anterior temporal and frontal responses.

**Figure 2:**
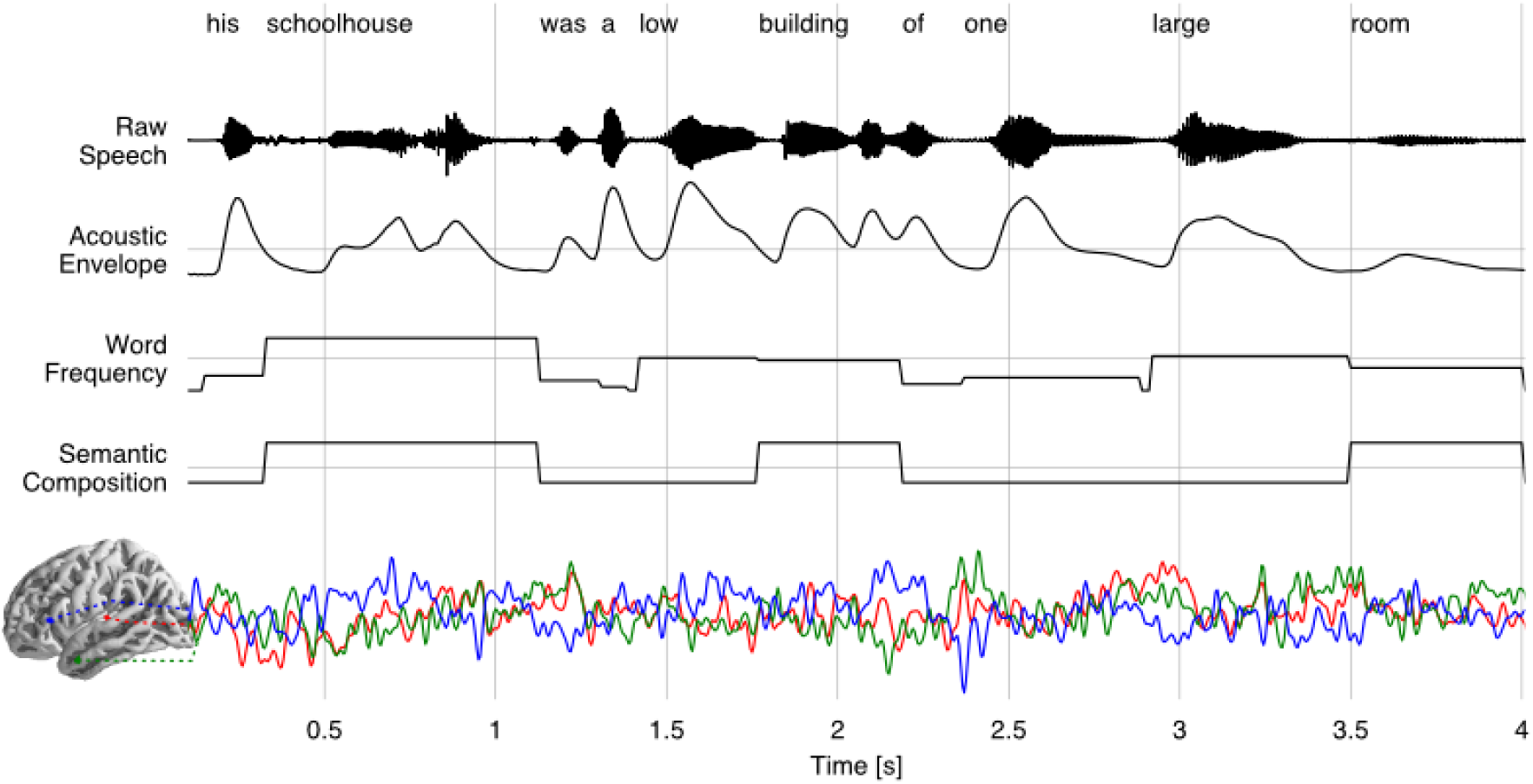
Stimulus coding for kernel estimation. Illustration of the first 4 seconds of one of the two speech stimuli. The text at the top indicates the transcript; the next four lines show the raw acoustic waveform data and the three continuously coded predictor variables. The bottom illustrates the source localized MEG data from three arbitrary source dipoles from a representative participant, averaged across the three presentations of the stimulus. The analysis modeled the brain signal at each source dipole based on the three predictor variables using convolution with a kernel of 1 second length.

## 2 Methods

### 2.1 Testing dataset

We analyzed a subset of the data described in detail by Presacco and colleagues (Presacco, Simon, & Anderson, 2016). In brief, 17 adults (aged 18-27 years) recruited from the Maryland and Washington, D.C. areas listened to one-minute long segments of an audiobook recording of *The Legend of Sleepy Hollow* by Washington Irving (https://librivox.org/the-legend-of-sleepy-hollow-by-washington-irving/), narrated by a male speaker, and sampled at 44,100 Hz. Audio segments were modified to remove pauses longer than 300 ms. Stimuli were delivered diotically through foam earphones inserted into the ear canal at ˜70 dB sound pressure level, with a sound delivery system equalized for an approximately flat transfer function from 40 to 3,000 Hz. Each segment was repeated three times. While the recording sessions included conditions with two-speaker audio, for the present analysis, only two one-minute long segments of single speaker audio in quiet were used, for a total of 6 minutes of data from each participant. To maximize attention to the stimuli, participants were asked before presentation of each segment to silently count the number of times a specific word or name was mentioned.

Handedness of the participants was assessed with the Edinburgh handedness scale (Oldfield, 1971). The scale measures a lateralization quotient, which can range from −1 (complete left-dominance) to 1 (complete right-dominance). Results indicated right-dominance in the majority of our sample, with 15 out of 17 participants having a lateralization quotient > 0. To exclude the possibility that the tests for lateralization of brain responses were biased by including lefthanders, these tests were repeated including only participants with a lateralization quotient of 0.5 or larger (n = 13). While this did not reveal any additional significant effects, lateralization of the early acoustic response become non-significant as reported in the appropriate section below.

### 2.2 Predictor variables

Stimulus variables were created reflecting three cognitive levels of speech processing: acoustic power, word frequency and semantic composition. For linear kernel estimation, predictor variables were sampled at the same rate as the dependent variable, i.e. the source localized MEG data, at 100 Hz.

#### 2.2.1 Envelope

An auditory spectrogram representation was generated for each stimulus using a model of the auditory periphery (Yang, Wang, & Shamma, 1992). The auditory spectrogram is a frequency by time matrix reflecting the representation of the acoustic signal in the brainstem. This representation was averaged across frequency bands to generate a univariate continuous predictor reflecting momentary acoustic power at each time point.

#### 2.2.2 Word frequency

Phonemes in the speech stimuli were labeled using the Gentle forced aligner (Ochshorn & Hawkins, 2016), and phoneme boundaries were manually adjusted using Praat (Boersma & Weenink, 2017). In the analysis reported here, only word boundaries were used. Logarithmic word frequencies (log_10_wf) were retrieved from the SUBTLEX database (Brysbaert & New, 2009). They were encoded into a continuous predictor with value 0 during silence, and 6.33 – log_10_wf for the duration of each word. This value was chosen to code infrequent words as high values, and very frequent words as low values; the highest log_10_wf entry in the database is 6.329.

#### 2.2.3 Semantic composition

A variable approximately tracking the amount of semantic composition across the speech stimulus was created by identifying all word groups corresponding to the semantic composition patterns identified by Westerlund et al. (2015): adjective-noun, adverb-verb, adverb-adjective, verb-noun, preposition-noun and determiner-noun pairs. The second word of each pair was marked. Simple articles (*the, a*) were ignored when identifying determiners because of their low semantic content. In patterns with multiple modifiers, the head word was marked in the same way; for example, in *a substantial Dutch farmer,* with two adjectives modifying the same noun, the noun *farmer* was marked. Words associated with semantic composition were coded as 1 for the duration of the word, all other time points as 0.

#### 2.2.4 Correlations between regressors

Time-point by time-point, both word-based predictor variables were only weakly correlated with the acoustic envelope predictor variable (word frequency: *r* = .08; semantic composition: *r* = .09). The correlation between the two word-based variables was larger (*r* = .39), owing to the fact that only content words were candidates for our semantic composition predictor, and content words tend to have lower frequencies than function words (the correlation is positive because lower frequency was coded with higher values).

### 2.3 MEG data acquisition and preprocessing

Before the experiment, each participant’s head shape was digitized with a Polhemus 3SPACE FASTRAK system, including 3 fiducial points and 5 marker positions. Five marker coils attached to the subject’s head at the position of the marker points were used to localize the head position relative to the MEG sensors at the beginning and end of the recording session. These head position records were also used to verify that participants’ head had not moved excessively over the course of the recording session. The average distance between pre- and post-experiment marker positions was 4.7 mm, with two participants exceeding 10 mm (10.6 mm and 14.8 mm).

During the recording, participants were resting in supine position, in a dimly lit magnetically shielded room. Data were acquired on a 157 axial gradiometer whole head MEG system (KIT, Kanazawa, Japan) at University of Maryland, College Park and recorded with an online 200 Hz low pass filter and a 60 Hz notch filter at a sampling rate of 1 kHz.

Data were pre-processed with mne-python 0.14 (Gramfort et al., 2013, 2014). Flat channels were automatically detected and excluded. Extraneous artifacts were removed using temporal signal space separation (Taulu & Simola, 2006), and data were band-pass filtered between 1 and 40 Hz with a zero-phase FIR filter (with mne-python’s default settings). The six 60 second long data epochs corresponding to stimulus presentation were extracted and downsampled to 100 Hz. At that point, channels were inspected based on their average correlation with neighboring sensors in the raw data; no channel had an average neighbor correlation coefficient below 0.3. For graphical display only, time series were upsampled to 500 Hz to minimize visual discretization artifacts.

### 2.4 Source localization

Head position measurements from the beginning and end of the MEG recording session were averaged and used to localize the subject’s head shape relative to the MEG sensors. The digitized head shape was used to coregister the ‘fsaverage’ brain model provided by FreeSurfer (Fischl, 2012) to each subject's head using uniform scaling, translation and rotation. A source space was defined on the white matter surface of the fsaverage model using four-fold icosahedral subdivision, with virtual current dipoles oriented perpendicular to the cortical surface. These were used to compute a cortically constrained distributed minimum 𝓁2-norm inverse operator (Dale & Sereno, 1993; Hämäläinen & Ilmoniemi, 1994) using a noise covariance estimated from empty room data and depth weighting parameter of 0.8 (Lin et al., 2006). Filtered MEG data were projected to source space. Dipoles lying on subcortical structures along the midline were removed, leaving a total of 4731 dipoles.

While the current study employed minimum 𝓁2-norm source estimates because they are widely used and well established, other approaches generating distributed source models could be substituted. One caveat concerns dipole orientations: Analysis of evoked responses often relies on source estimates that compute absolute dipole amplitude while discarding the direction of the current. This is appropriate when an increase in current is expected in response to a unique event. However, when analyzing continuous responses, where high pass filtering replaces baseline correction, a change of the sign in the current estimate is important information. Hence, directional (“fixed orientation”) source estimates, which preserve information about the dipole orientation, are preferable.

Since the current estimates were normalized for kernel estimation at each source dipole, corrections that weight data by source location, such as dSPM noise normalization (Dale et al., 2000), are not applicable.

### 2.5 Linear kernel estimation

The linear model relating *r_t_*, the response at time *t*, to the stimulus is given by

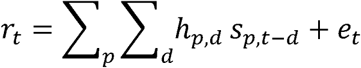

Where *S_p,t_* is the value of the stimulus variable for predictor *p* at time *t, h_p,d_* is the value of the kernel for predictor variable *p* at delay *d,* and *e_t_* is the prediction error (residual) at time *t*. The range of *d* determines which time points in the stimulus can influence the response at any time. For the results presented here, *d* ranged from 0.00 to 0.99 s, thus, for example, the predicted response at time *t*=20.0 was modeled as a weighted average of the values of the predictor variables at the time points from *t*=19.01 to *t*=20.0.

We used boosting with cross-validation and 𝓁1 error norm to estimate sparse response functions unbiased by the autocorrelation in the stimulus (for details see David, Mesgarani, & Shamma, 2007). The precise implementation is available in the Eelbrain source code (Brodbeck, 2017). Briefly, data were first divided into 10 equal contiguous parts along the time axis, and 9 parts were used as training data, the remaining part as test data. The boosting algorithm started with a response function of *h_0_* = 0 for its entire duration, which was iteratively modified at a single time point in increments of a constant Δ. Given a kernel and a stimulus array, the response is predicted by:

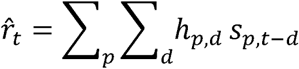

In each iteration, the training data was used to determine that element of *h* in which a change lead to the largest 𝓁1 error reduction; the resulting new kernel was then evaluated as to whether it reduced the error for the testing data. Iteration stopped when the error for the training data could not be reduced any further, or when the error for the testing data increased in two successive iterations. When iteration stopped, the kernel from the iteration with the smallest error on the testing data was retained.

This procedure was repeated 10 times, with each of the 10 data segments serving as test segment once. The 10 resulting kernels were averaged.

To make the iterative changes comparable across predictor variables, all stimulus as well as response variables were centered and normalized by dividing by the mean absolute value, and the change step was set to Δ= 0.005.

### 2.6 Incremental model comparison

The quality of the prediction of the signal at each virtual current dipole can be expressed by the correlation between the predicted and the actual response. This implies a straight-forward method for comparing the predictive power of different models across brain regions by comparing correlation maps.

To test whether adding a given predictor to the model leads to significant improvement, two models were fit for each subject: one with all three predictors of the full model, and a second one that was identical except that the predictor under investigation was temporally permuted to remove the relationship with the response data. For each of the one-minute long stimuli, the predictor was split in the middle and the first half, i.e. the first 30 seconds, of the stimulus was used to predict the neural response to the second half of the stimulus, and vice versa. This procedure removed the temporal relationship to the neural data while, while keeping the local temporal structure of the stimulus identical between the true and the permuted model. The Pearson correlation coefficients, expressing the fit between the predicted and the actual responses, were rescaled with Fisher’s z-transform, and one-tailed *t*-test were used to test whether the correctly aligned predictor improved the prediction of the neural data.

To control for multiple comparisons when testing for correlation coefficient differences at a large number of virtual current dipoles, we used nonparametric permutation tests (Nichols & Holmes, 2002; Maris & Oostenveld, 2007) based on the threshold-free cluster-enhancement algorithm (TFCE;Smith & Nichols, 2009). The precise implementation is available in the Eelbrain source code (Brodbeck, 2017). First, a *t*-value was computed for each virtual dipole based on the difference in correlation values across subjects. The resulting *t*-map was then processed with TFCE, an image processing algorithm that enhances contiguous areas of large values compared to isolated spikes, based on the assumption that meaningful differences have a larger spatial extent than noise. To determine a statistical distribution for the resulting TFCE values, we repeated the procedure in 10,000 random permutations of the data. In each permutation, condition labels were flipped for a randomly selected set of subjects, without sampling the same set of subjects twice (i.e., in each of the 10,000 permutations, the labels for at least one, and at most all, subjects were flipped). The *t*-test and TFCE were repeated for each permutation, and the largest value from the cluster-enhanced map was stored as an entry in the distribution. Thus, we computed a nonparametric distribution for the largest expected TFCE value under the null hypothesis. Any value in the original TFCE map that exceeded the 95^th^ percentile of the distribution was considered significant at the 5% level, corrected for multiple comparisons across the whole brain.

### 2.7 Evaluation of response functions

In addition to the model fit, the boosting algorithm results in an estimated response function at each virtual dipole for each predictor. These response functions contain information about the time course with which the information in different predictors affected different brain regions. Because boosting tends to result in temporally sparse response functions (cf. Figure 1), response functions for each subject were smoothed with a 50 ms (5 sample) Hamming window to make them more amenable to group analysis. Since the window was centered, distortions of peak latencies are not expected, but effects might appear slightly more temporally extended than they are in individuals’ responses.

Since directional current estimates were used, the expected mean of each source was 0. Response functions were thus tested with two-tailed one-sample *t*-tests against 0. Control for multiple comparisons across time and anatomical space was implemented with the same method as for model comparisons, except that the data had the additional dimension of time.

In MEG source localization, the signal at several thousand virtual current dipoles is estimated based on measurements at a much smaller number of sensors, in our case 157 axial gradiometers. Source localization accuracy is thus inherently limited; minimum 𝓁2-norm estimates tend to be spatially smeared. Since the minimum 𝓁2-norm inverse operator is a linear matrix operation, the source localization accuracy can be characterized by the point spread function (Hauk, Wakeman, & Henson, 2011). The point spread function describes the source estimate for a hypothetical point source, by projecting the activity in that one source dipole to the MEG sensors, and then applying the inverse solution to project the estimated magnetic fields back to the current source dipoles of the source space. Figure 3 illustrates the point spread function of the KIT 157 sensor MEG system and its influence on source localization at the group level. These exemplary plots illustrate that the spatial extent of the current estimates must be interpreted with care.

**Figure 3:**
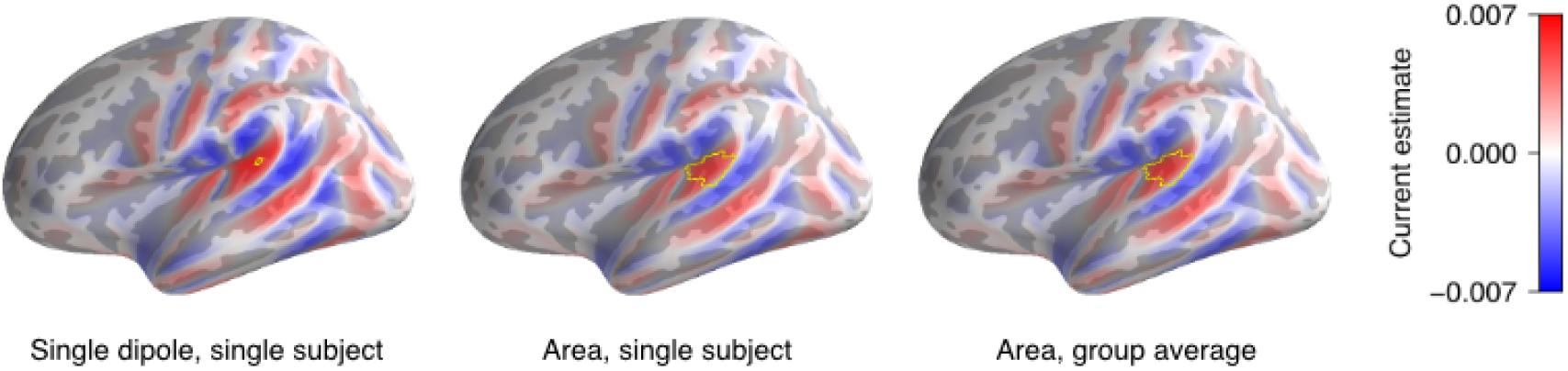
Point spread function. The point spread function is a theoretical estimate of the maximum spatial precision that can be expected in MEG source estimates. It is specific to a given MEG sensor configuration and head geometry; these plots are based on the specific details of the present study. Given that both forward operator and inverse operator are linear matrix operators, transforming data from source space to sensor space and back, forward and inverse operators can be combined to give the hypothetical source estimate for a single active dipole (left). Point spread functions can be locally summed to give the hypothetical source estimate for an area of active dipoles (middle). Finally, individual subjects' point spread functions can be combined for an estimate of the spatial accuracy of group results (right). In the plots above, active dipoles are circled in yellow. Source current was normalized so that the sum of all currents was 1 in each of the 3 plots. Forward and inverse solutions were taken from the 17 subjects whose data were analyzed.

Due to the spatial dispersion, anatomically separate sources only lead to cleanly separated source estimates if their spatial separation is large relative to the point spread function. Consequently, typical response function estimates may contain multiple, partially overlapping activations, making interpretation of raw plots of source space responses more difficult. A critical part of interpreting response functions thus consists in disentangling overlapping responses, and in determining which activations reflect true independent neural sources, and which merely reflect artefactual spatial dispersion from a genuine source to nearby areas. In order to facilitate this task, we tested two methods for identifying unique sources of variability in the response functions: hierarchical clustering, and sparse principle component analysis (sPCA). Both methods make use of the excellent temporal resolution of MEG to identify separable sources of variability in the time course. However, they do so using different constraints: Hierarchical clustering attempts to find a small number of average cluster time courses, and can use a spatial constraint to generate contiguous clusters. A downside is that current directionality (negative or positive current) has to be discarded to prevent the sign from dominating the cluster mean (compare with the striping in Figure 3). SPCA, on the other hand, accommodates current direction reversals through components with negative weights; however, it cannot impose a spatial contiguity constraint, leading to distributed and partially overlapping components. Because each method has advantages and disadvantages, we present both for comparison. Both methods were implemented with functions from scikit-learn 18.1 (Pedregosa et al., 2011).

#### 2.7.1 Hierarchical clustering

Hierarchical clustering (Ward, 1963) was used to group dipoles with similar time courses (for an application to fMRI data see Thirion, Varoquaux, Dohmatob, & Poline, 2014). First, response functions were masked at the 5% significance level (based on the spatio-temporal TFCE test described in Section 2.7) to restrict clustering to aspects of the responses that were reliable across subjects: Non-significant elements were set to zero, and dipoles that included no significant element at any time point were discarded. Because clustering was based on the mean time course in each cluster, the absolute values of all response functions were used for clustering. This was done to avoid distorting clusters based on anatomical features, which lead to source estimates with alternating sign across gyri and sulci due to alternating cortical surface orientation (compare Figure 3).

The clustering algorithm successively merged sources to minimize the sum squared error from cluster means, until a complete tree incorporated all sources. Links were constrained such that no direct links could be formed between sources further apart than 10 mm in 3-dimensional space. This distance criterion was chosen over geodesic adjacency to account for the fact that, due to the orientation constraint of the source estimates, sources could be similar at elements of adjacent gyri with corresponding orientation, with intervening elements with a different orientation. This is particularly relevant for auditory activity, which may “leak” from the superior temporal gyrus across the Sylvian fissure into adjacent parts of the inferior parietal and frontal cortices. The hierarchical tree was then traversed from the root until implementing the next branching would have reduced the sum squared error by less than 1% of the total sum squared.

Because clustering is based on the cluster mean, whereas source estimates have a smooth center-surround shape, this procedure frequently leads to spurious clusters that form low amplitude “halos” surrounding other, higher amplitude peaks. Since such halos are due to spatial dispersion and do not reflect effects of interest, it was desirable to remove them for visualization. Two methods were employed to detect halo clusters: First, clusters whose time course peak was more than one standard deviation below the mean were flagged for removal. Second, pairs of clusters with a time course correlation larger than 0.9 were flagged for closer examination; if one constituted a clear halo of the other, it was removed. If both clusters exhibited independent spatial centers, they were merged into one cluster (this occurred only once, for cluster S1_*cl*_, whose parts likely reflect the same underlying neural source, but were not connected by the clustering algorithm due to the large spatial separation across the Sylvian fissure). Only clusters that conformed to the following criteria were classified as halos and removed: 1) a large spatial extent, surrounding one or more other clusters rather than covering its own center 2) markedly lower amplitude than the cluster at its center 3) no distinct peaks, except at the time points of the peaks of the cluster at the center (and with lower amplitude).

#### 2.7.2 Sparse PCA

Sparse PCA finds spatial components that optimally reconstruct the data with an iterative procedure, adding a sparsity constraint through an 𝓁1 penalty on the components (Mairal, Bach, Ponce, & Sapiro, 2009). The iterative algorithm attempts to minimize, for a given number of components, the error function

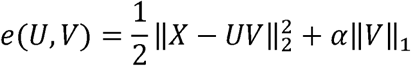

where X is the spatio-temporal response to be explained, U is the matrix of time courses and V is the matrix of sparse components. Analogously to the clustering procedure, response functions were masked by significance at the 5% level before submitting them to sPCA. The α parameter controlling sparsity was set to the largest (i.e., most sparse) power of 10 at which models still regularly converged: 10^−4^. The number of components was initialized with 1, and additional components were added until adding another component would have decreased the error by less than 1% of the total sum squared.

For visualization, sPCA components were normalized by setting the largest absolute value on each component map to 1, and component amplitude time courses were scaled appropriately. For display only, anatomical component maps were smoothed with a Gaussian with 5 mm standard deviation.

### 2.8 Test of lateralization

We tested for functional lateralization of responses by comparing response functions in the left and the right hemisphere. To perform a continuous spatio-temporal comparison, response functions had to be projected (“morphed” in FreeSurfer terminology) to a common hemisphere, i.e., data from one hemisphere had to be “mirrored” onto the other. While the fsaverage brain used for source estimation is slightly asymmetric, FreeSurfer also provides an exactly symmetrical brain, labelled “fsaverage_sym”, for the express purpose of hemispheric comparison (Greve et al., 2013). Because the fsaverage brain model is not precisely symmetric, projecting from one hemisphere to the other is not precise on the level of gyri and sulci (for example, the crown of a gyrus in one hemisphere might come to lie on the wall of the gyrus in the other). Thus, to avoid spurious differences due to current direction, response functions were first transformed to absolute values and Gaussian smoothing was applied with a full width half maximum of 10 mm. The resulting response functions from both hemispheres were then projected to the left hemisphere of the fsaverage_sym brain and used to compute left-right difference values at each source element, for each subject (projecting to the right instead of the left hemisphere did not substantively alter any results).

The left-right difference maps were masked to include only points in time and space at which the response function was significant in at least one hemisphere. For this, the original response functions were converted into a binary map at a threshold of *p* = .05 and projected to the left fsaverage_sym hemisphere. Since this resulted in some smoothing, the map was again binarized using a threshold of 0.5.

The resulting masked left-right difference maps were tested with two-tailed *t*-tests against 0, using the same permutation procedure with TFCE as for other tests to correct for multiple comparisons across space and time.

For visualization purposes, the resulting maps were again binarized at *p* = .05, and elements with negative differences were removed from the left hemisphere and projected to the right hemisphere. The resulting significance map covering both hemispheres was projected back to the fsaverage brain, and again thresholded at 0.5, resulting in a map of significant lateralization in space and time.

## 3 Results

The model fit was evaluated using the Pearson correlation between the actual and the predicted responses. Each of the three predictor variables was evaluated as to whether it significantly improved the model, by comparing the fit of the full model with the fit of a model in which this variable was deliberately misaligned, using the first half of the stimulus to predict the second half of the response and vice versa. Whole brain maps of the difference were tested for significance with one-tailed *t*-tests, correcting for multiple comparisons using permutation tests with threshold-free cluster enhancement (TFCE;Smith & Nichols, 2009).

Figure 4 shows the regions where each predictor had significant explanatory power. Results indicated highly significant contributions from each of the three predictors (all *p* < .001). The plots in Figure 4 are suggestive of localization differences, with semantic composition showing more anterior peaks than the acoustic envelope and word frequency. However, the large spatial extent of the effects, in particular of the acoustic envelope, also raises the strong possibility of leakage due to the spatial dispersion of MEG source estimates. While it may be that the response to the acoustic envelope is more distributed than the response to the other two variables, it is also quite possible that the acoustic representation in auditory cortex has a higher signal to noise ratio (SNR), and hence leads to spurious significant correlations at more distant sources due to spatial dispersion. This ambiguity also limits the usefulness of direct statistical comparisons of *r*-maps for testing hypotheses of localization differences between different predictors, because differences in SNR can obscure differences in localization.

**Figure 4:**
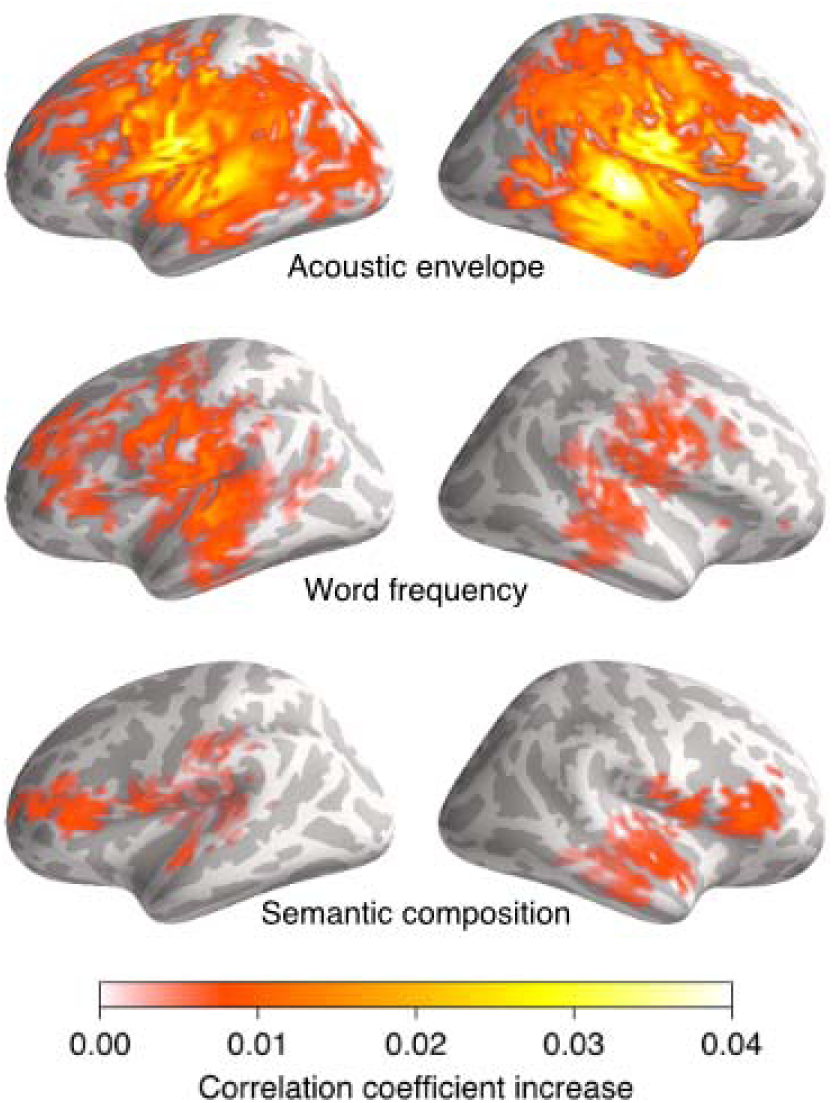
Significant model contributions. Each predictor variable was assessed for significant model contributions by comparing the fit of the full model to a model in which the predictor under investigation was temporally misaligned with the response. Each plot shows the difference in correlation-coefficient between the correct and the misaligned model at each dipole. Maps are thresholded by statistical significance, corrected for multiple comparisons across the whole brain, at p = .001 for the acoustic envelope and p = .05 for word-related predictors (the different thresholds account for the difference in signal to noise ratio in the neural representation of the predictors, and are used for graphical display purposes only).

In contrast to the model improvement maps, which condense all responses into a single anatomical map, the estimated response functions partition predictive contributions over time. Response functions thus have the potential to better show more nuanced distinctions between contributions from different brain regions via their processing latency differences. Furthermore, because all response functions were computed concurrently as a multidimensional kernel, the predictors were practically competing for explaining variance in the response. Consequently, each response function should reflect those components of the response that were best explained by the corresponding predictor, and exclude components that were better explained by other predictors in the model.

Figure 5 shows the response function to the acoustic envelope, masked by significance at the *p* = .05 level, and Figure 6 shows responses to the two word-related predictors. Response functions for each predictor were tested for regions with significant deviation from zero across anatomical space and response time with permutation tests, using TFCE. While response functions were estimated and tested for time delays from 0 to 1000 ms, they are shown in plots from 0 to 800 ms only because the last 200 ms were generally flat. The response functions exhibit some clearly identifiable peaks, which are localized more distinctly than the peaks of the correlation maps due to temporal separation of different peaks.

**Figure 5:**
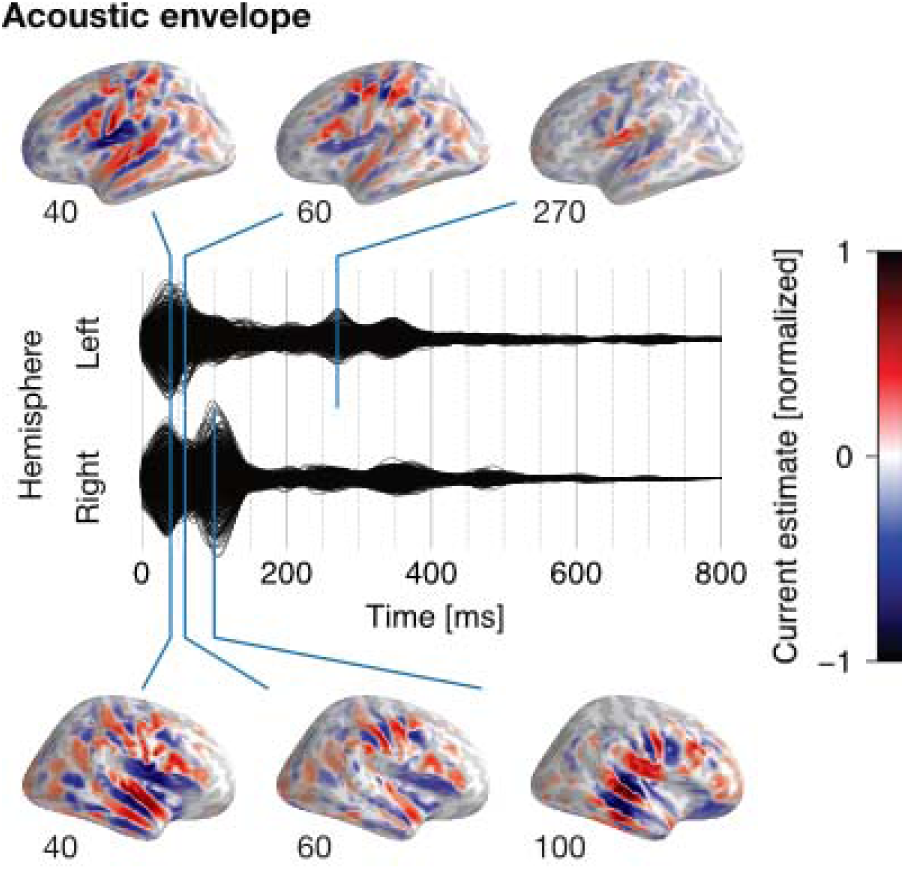
Acoustic envelope response function. Each black line reflects the response function at one virtual current dipole, averaged across subjects. Lines are separated by whether the corresponding dipole is part of the left or the right hemisphere. All values not significant at the 5% level, corrected across the whole brain, were set to zero. Anatomical plots show current distributions at visually obvious peaks, as well as peaks that emerged in the clustering analysis. Anatomical plots are rendered on the inflated surface of the fsaverage brain (for anatomical labels see Desikan et al., 2006). Numbers next to brain plots indicate time in ms.

**Figure 6:**
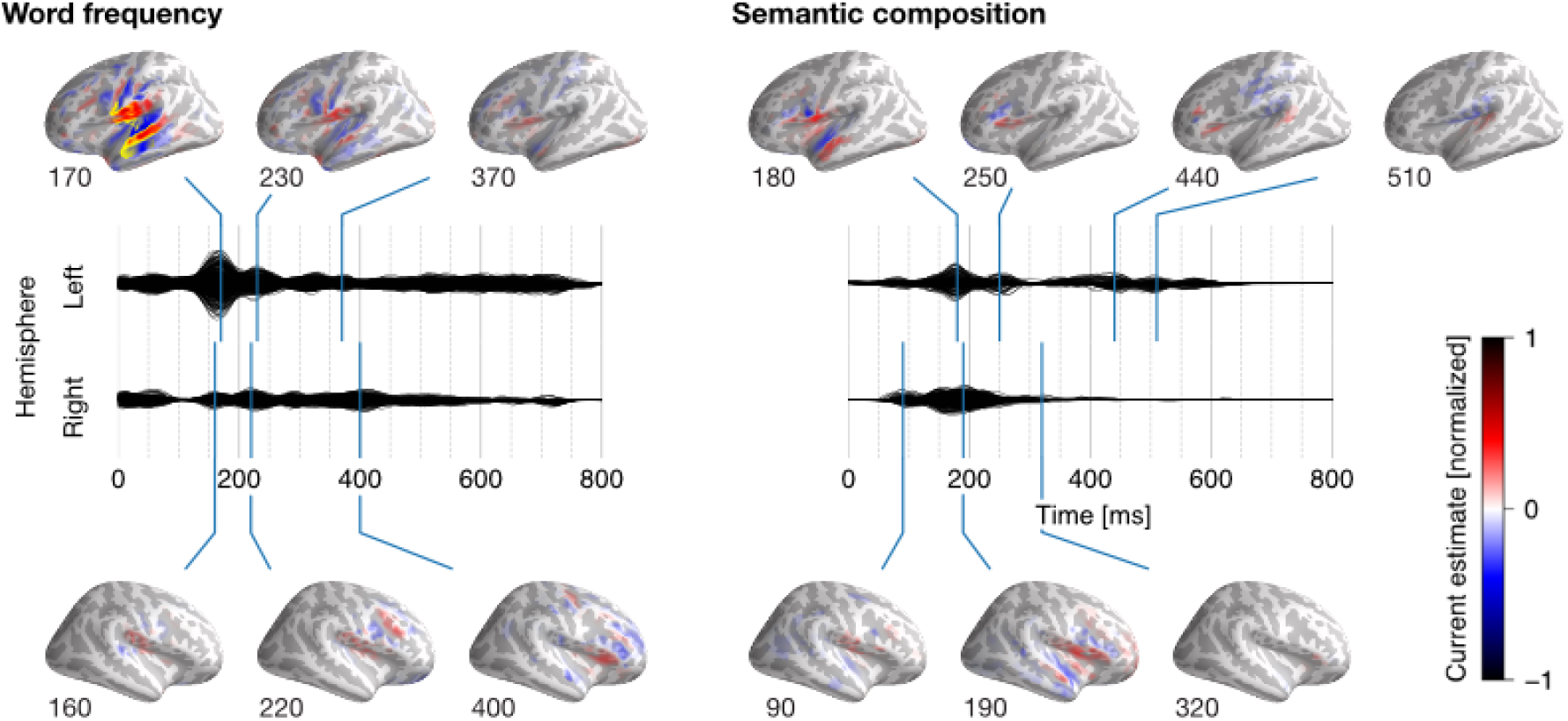
Word-related response functions. Plots are analogous to, and scale is identical with Figure 5. Areas of significant lateralization are indicated with yellow on the hemisphere with higher amplitude in the anatomical plots. The only plot with significant lateralization is the left hemisphere for word-freguency at 170 ms.

The acoustic envelope was associated with a first response peak around 40 ms, centered on auditory cortex bilaterally, and a second peak around 100 ms, localized slightly posterior to the first. Even though the second peak appears larger in the right hemisphere, the difference between hemispheres was not significant (superior temporal and Heschl’s gyrus, between 75 and 125 ms, *p* = .449). Around 60 ms, after the first response in auditory cortex, the response appeared to shift bilaterally to lateral Rolandic cortex, dorsal to auditory cortex, and to the inferior frontal gyrus. While this dissociation is harder to isolate in the raw response function, it was confirmed in the hierarchical clustering (c.f. clusters Al_*cl*_ and A2_*cl*_ in Figure 7) as well as the sPCA analysis (components A2_*pca*_ and A3_*pca*_ in Figure 8). These effects are clearly distinguishable from the auditory cortex response by their latency, and are unlikely to be due to spatial dispersion because the response in auditory cortex at 60 ms was much reduced. Thus, response functions were able to separate what looked like one large effect in the correlation maps presented above into multiple temporally distinct response components with clearly different neural sources.

**Figure 7:**
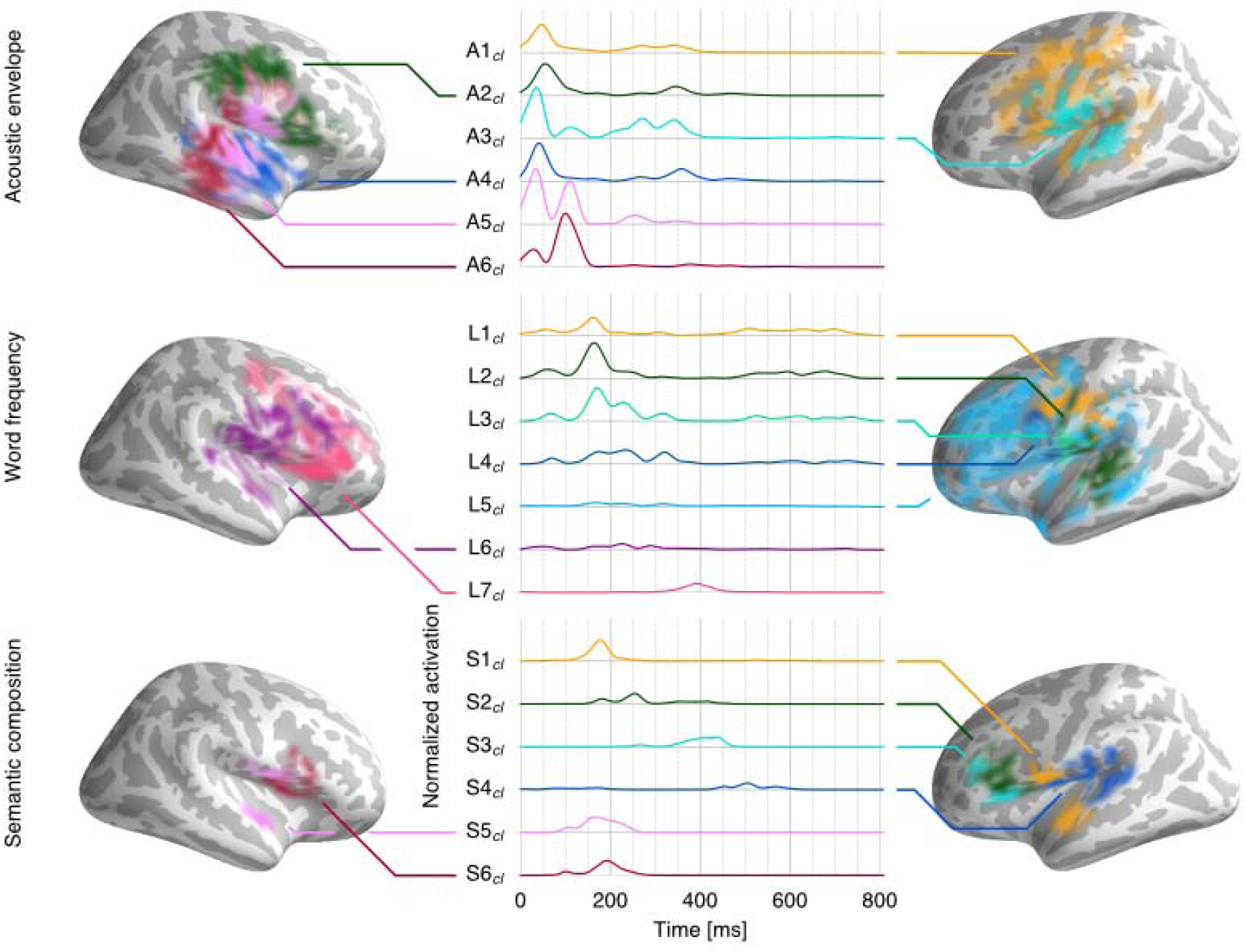
Responses grouped with hierarchical clustering. Each cluster is plotted with a color corresponding to the time course plot in the same color (with arbitrary numbering). The normalized activation (alpha) of the cluster reflects relative source amplitude within the cluster. The time course was computed as a weighted average of source time course, with weights determined by norm of each source. Clusters are labeled with a prefix corresponding to the predictor: (A) acoustic envelope, (L) lexical freguency, (S) semantic composition.

The lateralization test indicated a marginally stronger response in the anterior STG of the right hemisphere at 10 ms (*p* = .036). While the extent was small (the significant region encompassing only two source elements and one time point), numerically the difference extended throughout the M50 response (compare the 40 ms plots in Figure 5). This result could thus indicate a slightly stronger and/or earlier onset of an anterior component of the auditory response in the right compared to the left hemisphere; however, due to its marginal size we hesitate to interpret this effect further. When the test was repeated including only righthanded participants with a lateralization quotient of 0.5 or larger (n = 13), the lateralization test for the acoustic response lost significance (*p* = .098).

Word frequency was associated with a strongly left-lateralized response peak over auditory cortex around 170 ms. This peak was significantly larger in the left hemisphere than in the right hemisphere (*p* = .002). Areas with significant lateralization are shaded yellow in the anatomical plots of Figure 6. This response was followed by responses in the frontal cortex of both hemispheres.

The semantic composition predictor was associated with a progression of responses from anterior superior temporal gyrus to the lateral frontal lobes. The left hemisphere response exhibited clearer peaks, with a peak in the temporal lobe around 180 ms and an inferior frontal response peak around 250 ms, but hemispheric differences were not significant (*p* = .159). In addition, the left hemisphere response function exhibited late, weaker auditory cortex activation after 400 ms.

While response functions are more informative than model fit, it can be challenging to interpret them in terms of underlying neural sources due to the spatial dispersion, which can create a spatially complex pattern of results from a single neural source. However, source localization does not distort the time course of neural activity. We sought to utilize this fact to infer independent sources underlying the observed response functions based on the time-course of the responses. We tested two approaches for decomposing response functions into underlying sources based on their time course: hierarchical clustering and sparse principle component analysis (sPCA).

The first approach was based on hierarchical clustering of dipoles with similar time-course, while enforcing a realistic spatial layout by constraining possible groupings by anatomical proximity (Figure 7). This procedure identified 6 clusters in the acoustic envelope response function (after 2 halo artifacts were excluded). Clusters A3_*cl*_ in the left, and A4_*cl*_-A6_*cl*_ in the right hemisphere confirm the distinct localization of the two response peaks in auditory cortex. In addition, Al_*cl*_ and A2_*cl*_ identified a distinct response over Rolandic and inferior frontal areas of both hemispheres, with a slightly delayed peak compared to the early auditory cortex response. For word frequency, 7 clusters were identified (2 excluded halo artifacts). While the raw response function displayed in Figure 6 suggested one large peak in left auditory cortex, clusters L1_*cl*_−L4_*cl*_ suggest that this response is actually made up of distinct components, starting with a more posterior response in auditory (Ll_*cl*_) and possibly sensory-motor cortex (2_*cl*_), which is followed by a more anterior peak (L4_*cl*_); L3_*cl*_ might reflect a blend of L2_*cl*_ and L4_*cl*_. In addition, the clusters L5_*cl*_ −L7_*cl*_ draw attention to weaker but consistent frontal responses in both hemispheres. For semantic composition, 6 clusters (3 excluded halo artifacts; one cluster created from merging 2 others) were identified. In the left hemisphere, cluster S1_*cl*_ likely reflects an underlying source in the superior anterior temporal lobe with spatial dispersion across the Sylvian Fissure; S2_*cl*_ and S3_*cl*_ show a progression of activity to more anterior regions in the inferior frontal gyrus, while S4_*cl*_ might indicate later feedback to auditory regions. The right hemisphere showed a more homogeneous anterior temporal and inferior frontal response in clusters S5_*cl*_ and S6_*cl*_.

The second approach to isolating the sources underlying the response functions employed sparse principle component analysis (sPCA) to decompose response functions into a small number of spatially fixed patterns with time-varying amplitude. For the response to the acoustic envelope, sPCA isolated 4 components (Figure 8). In addition to reproducing the auditory cortex peak responses at 40 ms (Al_*pca*_) and 100 ms (A4_*pca*_), the results suggested that the intermediate response over the central sulcus can be divided into two components. A2_*pca*_ identified a slightly more dorsal peak around 50 ms, and A3_*pca*_ identified a slightly more ventral peak at 70 ms. Because the sPCA procedure did not impose any constraints on the spatial topography of the components, finding largely symmetrical components suggests that the effects were bilateral with very similar time course. The results for the lexical frequency response (Figure 9) confirmed the split of the left-dominant response into a main component over posterior STG (L1_*pca*_), and a more dorsal (L2_*pca*_) and a more anterior (L3_*pca*_) secondary component. This is interesting for the more dorsal L2_*pca*_ in particular, because components in sPCA are not formed based on amplitude but only based on the time course, thus, unlike in hierarchical clustering, weaker amplitude cannot be the sole explanation for a separate component. Rather, this result suggests the possibility that the response to lexical frequency is also associated with a component in left Rolandic cortex. A comparison of the primary response to lexical frequency, L1_*pca*_, with the primary acoustic response, A1_*pca*_, suggests closely aligned neural sources in the left hemisphere. Late frontal cortex responses with lower amplitude were also identified (L4_*pca*_ and L5_*pca*_). The 6 components identified in the semantic composition response suggest very similar conclusions as the hierarchical clustering (Figure 9). A single component centered on the superior anterior temporal lobe (S1_*pca*_) indicates that the clusters merged into S1_*cl*_ were indeed due to the same underlying source.

**Figure 8:**
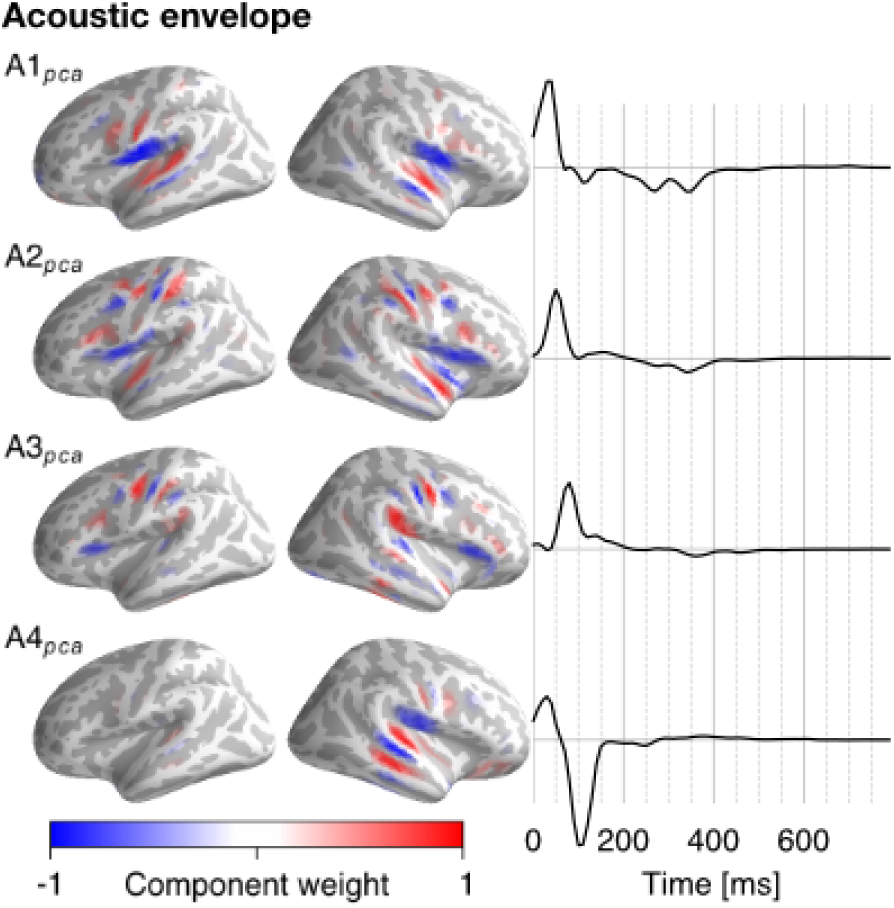
Acoustic response sparse PCA. Anatomical plots show the weights in the sPCA components, and time courses show the loading of the response function on the relevant component at each time point. All components are normalized such that the largest absolute value on each anatomical map is 1, and the time courses are shown to scale relative to each other. The sign of the components is inherently arbitrary, because simultaneously flipping the sign of a component and the corresponding time course leads to identical results; to make the components more comparable, the sign of overlapping components was manually adjusted to align the current direction in the area of overlap.

**Figure 9:**
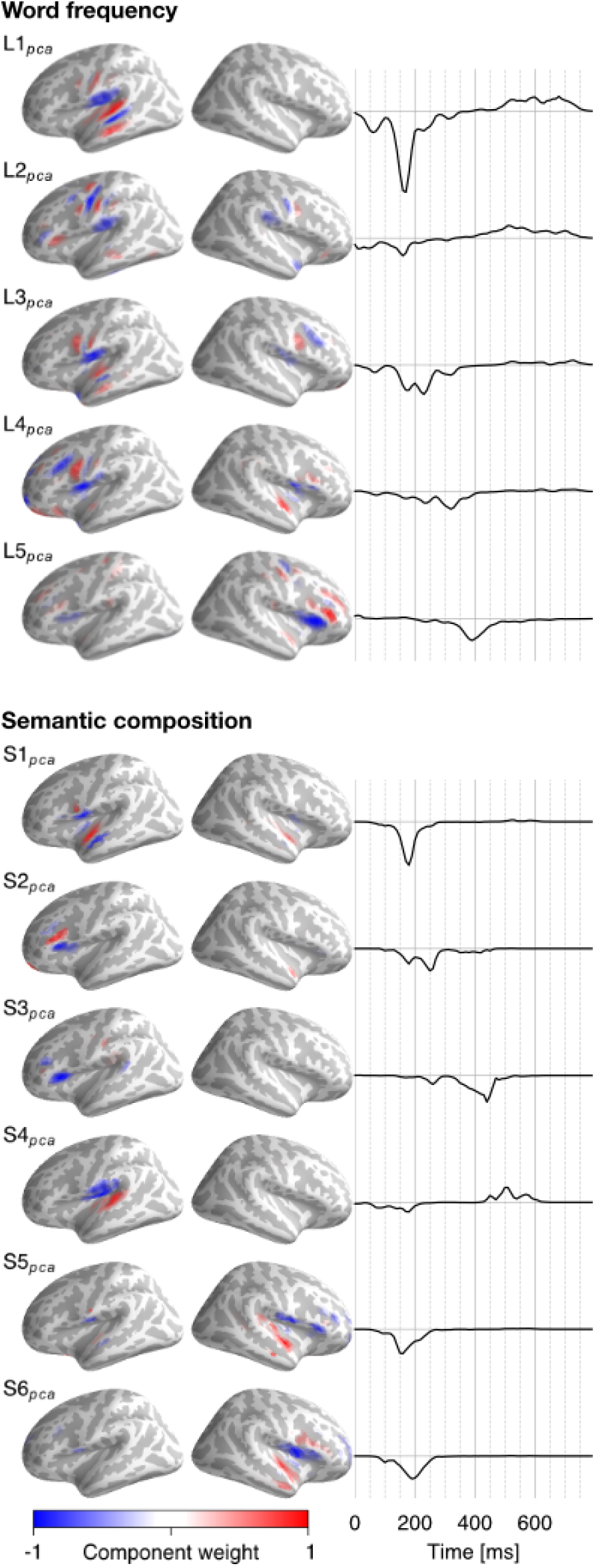
Word-related response sparse PCA. Details as in Figure 8, except that the response time courses were scaled by a factor of two.

## 4 Discussion

We described a procedure for combining linear kernel estimation with distributed MEG source localization to estimate the brain response to continuous stimuli as a function of delay time and anatomical space. We demonstrated the utility of this procedure by analyzing responses to continuous speech. Using just 6 minutes of MEG data per subject, we found reliable responses reflecting variables related to different cognitive levels of speech comprehension, including acoustic, lexical and semantic processing. Examination of the response functions revealed a detailed picture of the spatio-temporal evolution of cortical responses to continuous speech.

The spatial resolution of the estimated response functions is limited by the underlying inverse model, which infers current flow over a large surface area from measurements at a comparatively limited number of sensors (cf. Figure 3). One possible approach to this issue is thresholding results to emphasize peaks, which are more accurate than the extent of activation estimates (e.g. Hauk et al., 2011). However, thresholding responses composed of separate peaks with different signal strength may also hide peaks with lower amplitude. Here we illustrate a different approach, identifying effects with different underlying neural sources by separating sources of variation in the time course of activity. Hierarchical clustering and sPCA both allowed visualization of separate, more specific activation patterns, likely arising from different neural sources (e.g. Figure 8). Still, interpretation of the results should be guided by an awareness of the spatial resolution of the current estimates: The center of a given activation cluster can be assumed to be comparatively reliable, while the extent is likely exaggerated due to spreading of the minimum norm estimates.

### 4.1 Response functions to speech

Not surprisingly, we found a robust response to the acoustic envelope of the speech signal. This variable has been shown to be associated with brain signals measured with EEG (Lalor & Foxe, 2010) and MEG (Ahissar et al., 2001; Ding & Simon, 2012b), as well as intracranial measurements (Nourski et al., 2009; Mesgarani & Chang, 2012). Response functions to acoustic power were significant from the earliest time points. While the earliest responses could be due to the smoothing of the response functions, short latency responses to acoustic properties of speech are consistent with data suggesting that cortical responses to sounds occur within about 20 ms (Nourski et al., 2014; Parkkonen, Fujiki, & Mäkelä, 2009).

Previous studies have shown an earlier response component with a peak latency around 50 ms that follows acoustic power in the stimulus regardless of attention, and a later response component around 100 ms that reflects acoustic power in the attended speech stream rather than the raw acoustic stimulus (Ding & Simon, 2012b, 2012a, 2013; Akram, Simon, & Babadi, 2016). Our results suggested that the earlier response is in fact separable into two components, with a first peak around 40 ms, localized in auditory cortex, and a subsequent response within 10-30 ms over the central sulcus, dorsal to auditory cortex, and the IFG in both hemispheres. The location of this second component over the central sulcus is broadly consistent with the mouth region of the somatosensory homunculus (Nakamura et al., 1998). Motor cortex involvement in speech perception is predicted by the motor theory of speech perception (see Galantucci, Fowler, & Turvey, 2006) and has been demonstrated with meaningless syllables and single word stimuli (Pulvermüller et al., 2006; Schomers, Kirilina, Weigand, Bajbouj, & Pulvermüller, 2015; Wilson, Saygin, Sereno, & Iacoboni, 2004). Recent evidence suggests that this functional integration is supported by tight anatomical connections between Heschl's gyrus and primary motor and somatosensory cortex (Skipper & Hasson, 2017). Our result of rapid bilateral responses related to the speech envelope is compatible with a bilateral mechanism for a unified sensory-motor representation of speech (Cogan et al., 2014) through responses tied to acoustic, more than phonemic or motor, properties (Cheung, Hamilton, Johnson, & Chang, 2016). This system is contrasted with more abstract mappings between acoustic and motor representations thought to be left-lateralized (Hickok & Poeppel, 2004; Saur et al., 2008) and, as evidenced by patients with left-lateralized brain lesions, is probably not *necessary* for speech comprehension (Rogalsky, Love, Driscoll, Anderson, & Hickok, 2011). Our results thus suggest that involvement of bilateral motor regions in speech processing is not restricted to the somewhat unnatural discrete listening tasks frequently used in research, but occurs also during processing of natural connected speech, and with a short latency relative to the acoustic signal.

Results also revealed a consistent response associated with the lexical frequency of the words being processed. This response was localized primarily to the auditory cortex of the left hemisphere, followed by frontal modulations of lower amplitude. This is consistent with fMRI results on auditory story comprehension, which found significant association with word-frequency in left STG and IFG (Brennan et al., 2016) as well as weaker right-hemispheric activity (Brennan et al., 2012). A comparison of sPCA components A1_*pca*_ and L1_*pca*_ suggests that the strongest response to word frequency originated from a location only slightly ventral to the primary auditory response. Language-specific processing in early auditory areas is consistent with the observation of selective responses to speech sounds early in the cortical hierarchy in STG (Nourski et al., 2014; Hullett, Hamilton, Mesgarani, Schreiner, & Chang, 2016) and is consistent with models placing lexical processing of speech in a hierarchy between the STG and the middle temporal gyrus (e.g. Overath, McDermott, Zarate, & Poeppel, 2015). The lateralization test revealed significant lateralization of this response component towards the left hemisphere. This suggests that in contrast to acoustic processing, lexical processing, as indexed by lexical frequency, is lateralized to the language-dominant hemisphere.

Semantic composition was associated with temporal and frontal lobe activity. Previous MEG research, using visually presented, strictly controlled minimal phrases, suggested that activation associated with semantic composition localizes most consistently to the anterior temporal lobe (Bemis & Pylkkänen, 2011; Westerlund et al., 2015). A study that compared responses to spoken and written two-word stimuli also found activity in a superior posterior temporal region comparable to our sPCA component S4_*pca*_ (Bemis & Pylkkänen, 2012). Lateral prefrontal cortex activation was not described in these studies. One possible explanation for this divergence is that the increased demand imposed by continuous speech leads to more extensive brain involvement for the same process. However, semantic composition in natural speech is also correlated with other structural properties of language, which have been associated with lateral prefrontal activation (Brennan et al., 2016; Nelson et al., 2017). While our stimuli do not provide enough variation to distinguish between different related variables, our demonstration that MEG is sensitive to these variables in continuous speech opens up new possibilities to disentangle contributions from different semantic and structural variables.

Finally, response functions allow comparisons of the time course of processing of different variables. Precise comparison between the acoustic envelope and the word-related predictors is difficult due to the temporal nature of the respective variables. Acoustic power was coded as momentary acoustic power with millisecond resolution, while words were coded as events that could be temporally extended over hundreds of milliseconds. Indeed, this is not just a matter of coding, but also of the time scale at which the information unfolds, with words reflecting integration of information over a larger time interval. Thus, while the acoustic power is clearly associated with an earlier response than word properties, direct comparison of peak latencies is difficult. On the other hand, the two word-related predictors were coded with the same temporal structure and can be directly compared. The main response to semantically composable words in the anterior temporal lobe peaked around 180 ms, only 10-20 ms after the main response to lexical frequency. This is consistent with the observations in two-word studies that composition-related activity can have a short latency, peaking 225 to 250 ms after visual word onset (Bemis & Pylkkänen, 2011, 2012; Westerlund et al., 2015). Together, these observations support the notion that lexical information is integrated quickly with the information that is already available from the preceding speech signal.

More generally, previous research suggests that low-frequency phase-locked brain activity is related to higher levels of language processing, consistent with higher level information occurring at slower rates (e.g. Ding, Melloni, Zhang, Tian, & Poeppel, 2015). In the present analysis, this is reflected in the predictor variables for word frequency and semantic composition, which are both dominated by temporal variations in the delta band (1-3 Hz). However, the linear filter model implies that a predictor cannot predict brain activity at frequencies it does not model; While the present results are thus consistent with low frequency phase locked activity in higher level language comprehension, this is a consequence of the model, and does not preclude the possibility of a cognitive process that could be modeled at higher frequencies. A challenge for future work will be modeling predictors for different aspects of the comprehension processes more accurately.

### 4.2 Using source localization with linear kernel estimation

Our results confirm the viability of combining source localization with linear kernel estimation to estimate brain responses to continuous stimuli. Significant contributions from different stimulus variables could be identified, and response functions provided more details on the anatomical and temporal properties of the brain’s response.

The present analysis described neural response functions by testing for responses that were significantly non-zero across participants. While this is useful for demonstrating that brain activity tracks a given stimulus variable at all, and for determining which brain regions are involved in processes related to this variable, more fine-grained analyses will be possible by comparing response functions to the same variable under different experimental conditions. Such statistical analysis would be a straightforward extension of the approach shown here, replacing the one-sample *t*-tests used for analyzing the response functions with two sample *t*-tests or ANOVAs. The present analysis suggests that robust response functions can be estimated from just 6 minutes of MEG data per subject, demonstrating that future experiments could estimate response functions for multiple experimental conditions.

Robust responses were discovered for predictor variables reflecting not only acoustic properties of the speech stimulus, but also lexical and semantic properties, attesting to the possibility of studying not only sensory, but also cognitive processing of continuous stimuli. Computing response functions for source localized data allowed us to separate the brain responses associated with different properties of the same speech stimulus anatomically. In addition, response functions revealed dynamics over the course of the response to each variable, with different brain regions responding at different latencies. Based on the fact that the temporal resolution of MEG source estimates exceeds their spatial resolution, it should be possible to identify different neural sources by the unique sources of variation over the time courses at different dipoles. We tested two such methods, hierarchical clustering and sPCA, with largely convergent results. On the whole, though hierarchical clustering leads to simpler visualization (see Figure 7), sPCA has the advantage of preserving current direction and allowing for spatially overlapping components. As a consequence, sPCA is able to separate underlying sources more cleanly, and is not susceptible to halos and blended clusters. For example, sPCA modeled the M50/M100 distinction as two overlapping components (Al_*pca*_ and A4_*pca*_), whereas hierarchical clustering resulted in three clusters, with an additional cluster for the region of overlap blending both components (A4_*cl*_-A6_*cl*_).

While the correlation between the acoustic predictor variable and the lexical and semantic variables was relatively low (r ≤ .09), the correlation between word frequency and semantic composition was higher (r = .39). This underlines the importance of modeling contributions from different predictors together, rather than independently. The present approach using boosting addresses this issue in two ways: First, one multidimensional kernel is estimated to predict the response from all predictors simultaneously, i.e., the predictors compete for explaining variance in the response. And second, by fitting a permuted baseline model for each predictor, we specifically test that adding the predictor improves the model after accounting for the other predictors of the full model.

The ability to detect temporally and anatomically distinct response components offers new possibilities for future research. Often, distinct response components correspond to different cognitive processing steps. For example, two response components to the speech envelope with peaks around 50 and 100 ms differ in their sensitivity to attention, suggesting that only the latter is sensitive to top-down attentional modulation and thus reflects a more invariant auditory object representation (Ding & Simon, 2012a). Thus, the ability to distinguish response components is instrumental in delineating cognitive processing stages.

For analyzing response to continuous stimuli this technique complements fMRI, which can localize slower hemodynamic changes with high spatial accuracy, but does not have the temporal resolution of MEG critical for rapidly evolving processes like language comprehension. For example, a study that assessed similar variables with fMRI (Brennan et al., 2012) sampled neural data and predictors at 0.5 Hz. The hemodynamic response was directly compared with predictor variables convolved with the hemodynamics response function, without modeling dynamic response functions for neural responses. While fMRI thus allowed more accurate anatomical localization, it did not allow comparisons involving the temporal evolution of the neural responses as were revealed by our results. For example, while fMRI could localize the effects of word frequency to the left temporal and frontal cortices (Brennan et al., 2016), our results detected a temporal progression, with temporal lobe responses preceding frontal lobe responses.

### 4.3 Limitations

A major limitation of distributed minimum norm estimates of MEG employed here is the comparably low spatial accuracy. Ultimately, the actual accuracy depends on a variety of factors, from the MEG system used to the choice of inverse solution, and is not uniform across the brain (see e.g. Hauk et al., 2011). This issue directly affects the analyses presented here, and results should be interpreted accordingly. The issue of spatial dispersion can be partly addressed by using the estimated current time course to group dipoles that are likely to reflect the same underlying neural source. We showed that two such methods, clustering and sPCA, provided useful summaries of the response functions. A comparison of sPCA components with the point spread function (Figure 3) suggests that many sPCA components could potentially be the result of a single localized neural source. In sum, while informative at larger scales, localization results should be interpreted with care at a scale below a few tens of millimeters.

A related limitation specific to the present study is the substitution of scaled average brains for source reconstruction. Structural MRIs, when available, would allow source estimates to be computed on more accurate, subject-specific anatomical models.

Finally, a number of specific data processing methods used in the current analysis could easily be exchanged for alternatives. In particular, the present analysis uses boosting to estimate response functions; other methods, such as ridge regression (Lalor et al., 2006), would constitute viable alternatives and might even be better suited for different designs. Similarly, the present analysis was based on assuming a linear filter model, but it could be extended to nonlinear filters to test more advanced hypotheses.

## 5 Conclusion

We demonstrated that linear kernel estimation can be combined with distributed minimum norm source estimates to map brain response to continuous speech in time and anatomical space. While we developed and tested this technique for studying speech processing, it is applicable to other continuous stimuli. Kernels can be estimated with multiple predictor variables competing for explanatory power, which makes it possible to model responses to suspected covariates and test whether variables of interest explain variance in the responses above and beyond the covariates. This makes it amenable to investigate a range of sensory and cognitive processes with more natural stimuli than hitherto possible.

## Acknowledgements

This work was supported by the National Institutes of Health [grant number R01-DC-014085].

